# Correcting for experiment-specific variability in expression compendia can remove underlying signals

**DOI:** 10.1101/2020.05.03.066597

**Authors:** Alexandra J. Lee, YoSon Park, Georgia Doing, Deborah A. Hogan, Casey S. Greene

## Abstract

**Motivation:** In the last two decades, scientists working in different labs have assayed gene expression from millions of samples. These experiments can be combined into compendia and analyzed collectively to extract novel biological patterns. Technical variability, sometimes referred to as batch effects, may result from combining samples collected and processed at different times and in different settings. Such variability may distort our ability to interpret and extract true underlying biological patterns. As more multi-experiment, integrative analysis methods are developed and available data collections increase in size, it is crucial to determine how technical variability affect our ability to detect desired patterns when many experiments are combined

**Objective:** We sought to determine the extent to which an underlying signal was masked by technical variability by simulating compendia comprised of data aggregated across multiple experiments.

**Method:** We developed a generative multi-layer neural network to simulate compendia of gene expression experiments from large-scale microbial and human datasets. We compared simulated compendia before and after introducing varying numbers of sources of undesired variability.

**Results:** We found that the signal from a baseline compendium was obscured when the number of added sources of variability was small. Perhaps as expected, applying statistical correction methods rescued the underlying signal in these cases. As the number of sources of variability increased, surprisingly, we observed that detecting the original signal became increasingly easier even without correction. In fact, applying statistical correction methods reduced our power to detect the underlying signal.

**Conclusion:** When combining a modest number of experiments, it is best to correct for experiment-specific noise. However, when many experiments are combined, statistical correction reduces one’s ability to extract underlying patterns.

## Introduction

For the last two decades, unprecedented amounts of transcriptome-wide gene expression profiling data have been generated, most of which are shared in public platforms for the research community.^1^ Researchers are now combining samples across different experiments to form compendia, and analyzing these compendia is revealing new biology.^2–6^ It is well-understood that technical sources of variability pervade large-scale data analysis such as transcriptome-wide expression profiling studies.^7–10^ Numerous methods have been designed to correct for various types of effects.^7,11,12^ Despite the prevalence of technical sources of variability, researchers have successfully extracted biological patterns from multi-experiment compendia without applying correction methods.^2–5,13^ We sought to determine the basis of these seemingly contradictory results by examining the extent to which underlying statistical structure can be extracted from compendium-style datasets in the presence of sources of undesired variability.

A number of methods have been developed to simulate transcriptome-wide expression experiments.^14–17^ However, simulating a compendium of many experiments with existing approaches would require defining a statistical model that describes the process by which researchers design and carry out experiments, which is likely to be very challenging. Instead, we developed an approach to simulate compendia by sampling from the low-dimensional representation produced by multi-layer generative neural networks trained on gene expression data from an existing compendium. This allowed us to simulate gene expression experiments that mimic real experimental configurations. We combined these experiments to create compendia.

Using this simulation approach, we studied how adding varying amounts of experiment-specific noise affects the statistical structure of gene expression compendia and our ability to detect underlying patterns. This topic is becoming pressing as more large-scale expression compendia become available. We found that prior reports of pervasive technical noise and analyses that succeed without correcting for it are, in fact, consistent. In settings with relatively few experiment-specific sources of undesired variation, the added noise substantially alters the structure of the data. In these settings, statistical correction produces a data representation that better captures the original variability in the data. On the other hand, when the number of experiment-specific sources of undesired variability becomes large, attempting to correct for these sources does more harm than good.

## Results

We characterized publicly available data compendia using refine.bio^18^, a meta-repository that integrates data from multiple different repositories. We found that an average experiment contained hundreds to thousands of samples in most widely studied organisms (Table 1). These samples were derived from hundreds to thousands of experiments, and the most common experimental designs had relatively few samples (medians from 5-12). We compared these to two readily available compendia, recount2 and one for *P. aeruginosa*, that have been used for compendium-wide analyses.^2,3,6^ The compendia that have been successfully used in prior work^2,3,6^ have similar median numbers of samples per experiment (recount2 = 4, *P. aeruginosa* = 6) to the current publicly available data.

**Table 1:** Public data usually have only a modest number of samples per experiment, though in aggregate many samples are available. Statistics for the 10 largest transcriptomic compendia found in refine.bio, which is a meta-repository containing publicly available expression data from the Sequence Read Archive (SRA)^19^, Gene Expression Omnibus (GEO)^20^ and ArrayExpress^21^.

|  | No. experiments | Median no. samples | Total no. samples |
| --- | --- | --- | --- |
| HOMO SAPIENS | 15440 | 12 | 571862 |
| MUS MUSCULUS | 13224 | 10 | 296829 |
| ARABIDOPSIS THALIANA | 1627 | 9 | 24855 |
| RATTUS NORVEGICUS | 1368 | 12 | 38530 |
| DROSOPHILA MELANOGASTER | 853 | 9 | 17836 |
| SACCHAROMYCES CEREVISIAE | 627 | 12 | 12972 |
| DANIO RERIO | 546 | 9.5 | 28518 |
| CAENORHABDITIS ELEGANS | 375 | 10 | 7953 |
| SUS SCROFA | 280 | 12 | 6063 |
| ZEA MAYS | 274 | 5 | 3458 |

### Constructing a generative model for gene expression samples

We developed an approach to simulate new gene expression compendia using generative multi-layer neural networks. Specifically, we trained a variational autoencoder (VAE)^22^, which was comprised of an encoder and decoder neural network. The encoder neural network compressed the input data through two layers into a low-dimensional representation and the decoder neural network expanded the dimensionality back to the original input size. The VAE learned a low-dimensional representation that can reconstruct the original input data. Simultaneously, the VAE optimized the lowest dimensional representation to follow a normal distribution (Figure 1A). This normal distribution constraint, distinguishes VAE’s from other types of autoencoders and allowed us to generate variations of the input data by sampling from a continuous latent space.^22^

**Figure 1.**
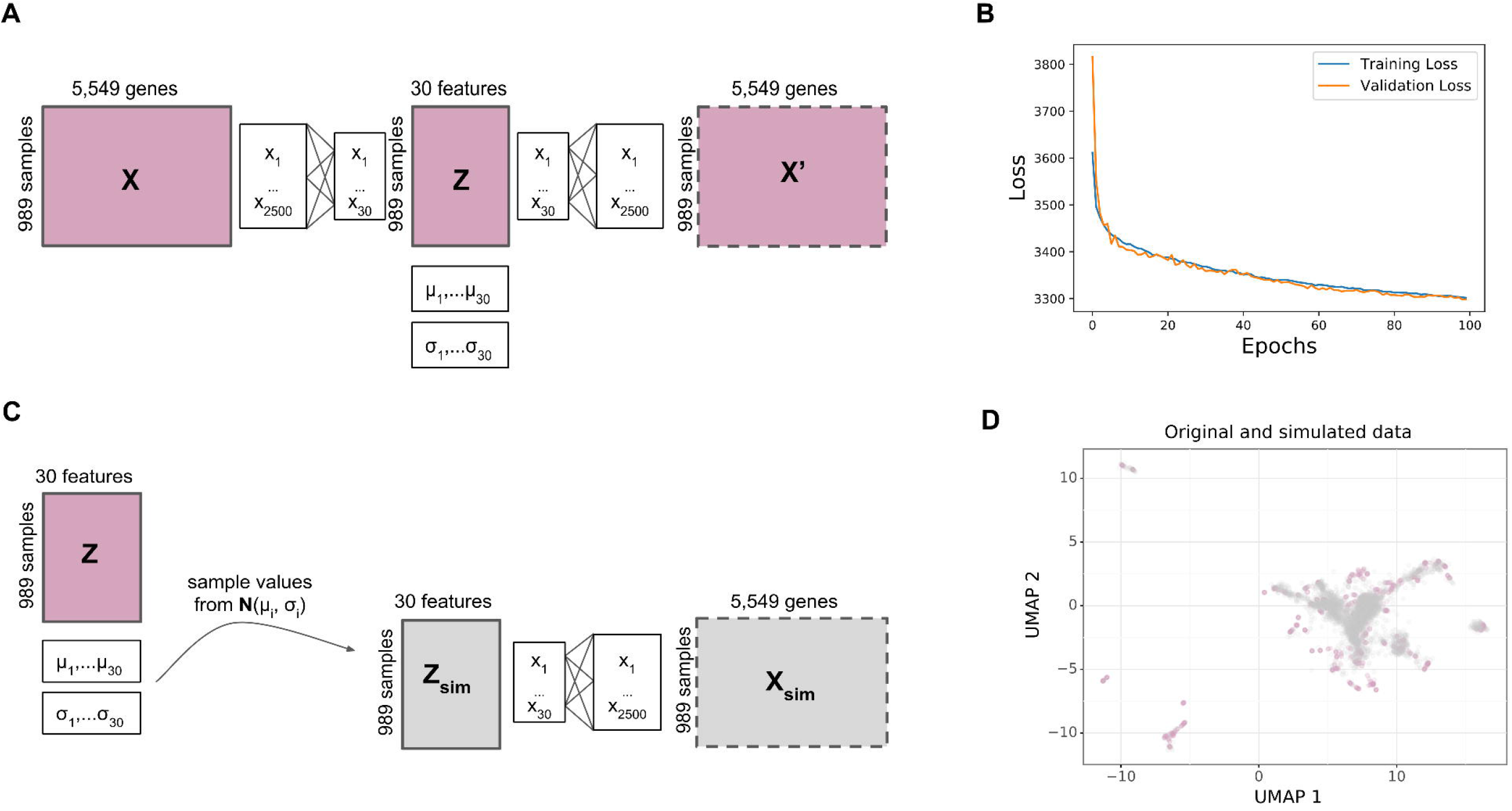
Simulating gene expression data using VAE. A) Architecture of the VAE, where the input data gets compressed into intermediate layer of 2500 features and then into a hidden layer of 30 latent features. Each latent feature follows a normal distribution with mean μ and variance σ. The input dimensions of the Pseudomonas dataset are shown here as an example (989 samples, 5549 genes). The same architecture is used to train the recount2 dataset except the input has 896 samples and 58,037 genes. B) Validation loss plotted per epoch during training using the P. aeruginosa compendium. C) Workflow to simulate gene expression samples from a compendium model, where new samples are generated by sampling from the latent space distribution. D) UMAP projection of P. aeruginosa gene expression data from the real dataset (pink) and the simulated compendium using the workflow in C (grey).

We trained VAEs for each dataset (recount2 and *P. aeruginosa*). We evaluated the training and validation set loss at each epoch, which stabilized after roughly 100 epochs (Figure 1B). We observed a similar stabilization after 40 epochs for recount2 (Figure S1A). We simulated new genome-wide gene expression data by sampling from the latent space of the VAE using a normal distribution (Figure 1C). We used UMAP^23^ to visualize the structure of the original and data and found that the simulated data generally fell near original data for both compendia (Figure 1D; Figure S1B).

### Simulating gene expression compendia with synthetic samples

We designed a simulation study to assess the extent to which artifactual noise associated with individual partitions of a large compendium affects the structure of the overall compendium. Our simulation is akin to asking: if different labs performing transcriptome-wide experiments randomly sampled from the available set of possible conditions, to what extent would experiment-specific biases dominate the signal of the data. We simulated new compendia, randomly divided the samples into partitions, and then added noise to each partition, and compared the simulated compendia with added noise to the unpartitioned one (Figure 2A). Each partition represents groups of samples with shared experiment-specific noise. We evaluated the similarity before and after applying an algorithm designed to correct for technical noise in each partition – given the linear noise added we used limma^24^ to correct.

**Figure 2.**
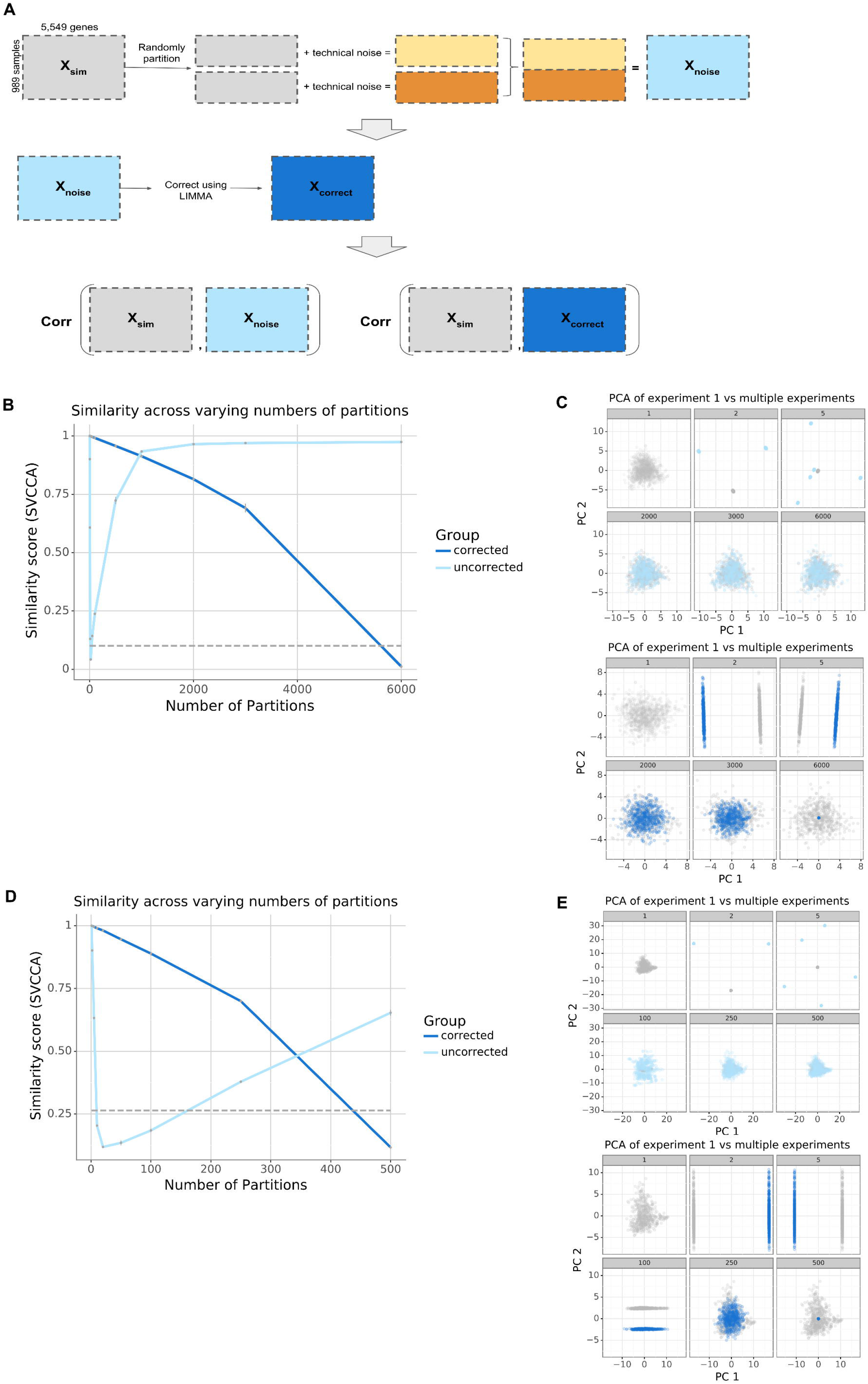
Results of simulating compendia. A) workflow describing how experiment-specific noise was added to the simulated compendia and how the simulated compendia were evaluated for similarity compared to the original input compendia. B,D) SVCCA curve measuring the similarity between a compendia without noise versus a compendium with noise (light blue), compendium with noise corrected for (dark blue). As a negative control, we used the similarity between the gene expression pattern of the simulated data with a single partition compared with the simulated data that has been permuted to destroy any meaningful structure in the data. C,E) Subsampled gene expression data (500 samples per compendia) projected onto the first two principal components showing the overlap in structure between the compendia without noise (gray) versus the compendia with noise (light blue), compendia with noise corrected for (dark blue).

We performed a study with this design using the VAE trained from the *P. aeruginosa* compendium for 2 to 6,000 partitions. We found that adding technical noise to partitions always reduced the similarity between the simulated data without partitions and the partitioned data. However, the nature of the change in similarity differed substantially between the partitioned sets before and after the correction step (Figure 2B). With the correction step, similarity dropped throughout the range of the study, eventually reaching the same level as the permuted data. Without the correction step, similarity dropped immediately to near the permuted level and then recovered throughout the rest of the tested range. Examining simulated data on the top 2 principle components from the original data with the corrected and uncorrected data at various numbers of partitions revealed that the correction step removes both wanted and unwanted variability, eventually removing all variability in the data (Figure 2C). Without correction, the data were initially dramatically transformed; however, as the number of partitions grows the effect on the structure of the data was diminished.

To determine whether or not this was a more general property of such compendia, we repeated the same simulation study using a VAE trained on a recount2 compendium. recount2 is a compendium comprised of human RNA-seq samples, so it is generated using a different technology and consists of assays of a very different organism. Results with recount2 mirrored our findings with the *P. aeruginosa* compendium. The correction step initially retained more similarity, but performance crossed over and by the end of the study the uncorrected data were more similar to the unpartitioned simulated compendium (Figure 2D). Examining the top principle components again revealed that correction better retained the structure of the original data with few partitions, but with many partitions the structure was better retained without correction (Figure 2E). We observed the same trends when we varied the magnitude of the noise added (Figure S2) or used a different noise correction method, such as COMBAT (Figure S3).

### Constructing a generative model for gene expression experiments

We randomly selected samples from the range of all possible samples in the compendium. For the next simulation, we developed an approach that could simulate realistic experimental structure. This next simulation added another level of complexity to the model, by simulating experiments as opposed to samples, in order to make the simulated compendia more representative of true expression data. The technique that we developed uses the same underlying approach of sampling from a VAE. However, in this case we randomly selected a template experiment and a vector that would move that template experiment to a new location in the gene expression space (Figure 3A). The simulation preserved the relationship between samples within the template experiment while also shifting the activity of the samples in the latent space (Figure 3B). Intuitively, this process maintained the relationship between samples but changed the underlying perturbation. We used this process to generate compendia of new gene expression experiments. We exampled how the original samples in an experiment (E-GEOD-51409) and a simulated experiment generated using E-GEOD-51409 as a template have consistent clustering of samples (Figure 3C original and experiment level simulated experiment).^25^ However the genes that were differentially expressed were different between the two datasets. This demonstrated that the perturbation intensity and experimental design were relatively consistent in gene expression space, even though the nature of the perturbation differed. The simulated experiment had a lower variance compared to the original dataset due to the normality assumption made by the VAE, which compresses the latent space data representation.^22^ However, in general, the clustering of samples is conserved between the simulated and original experiments, as observed in the additional template experiments with more complex structures (Figure S4).

**Figure 3.**
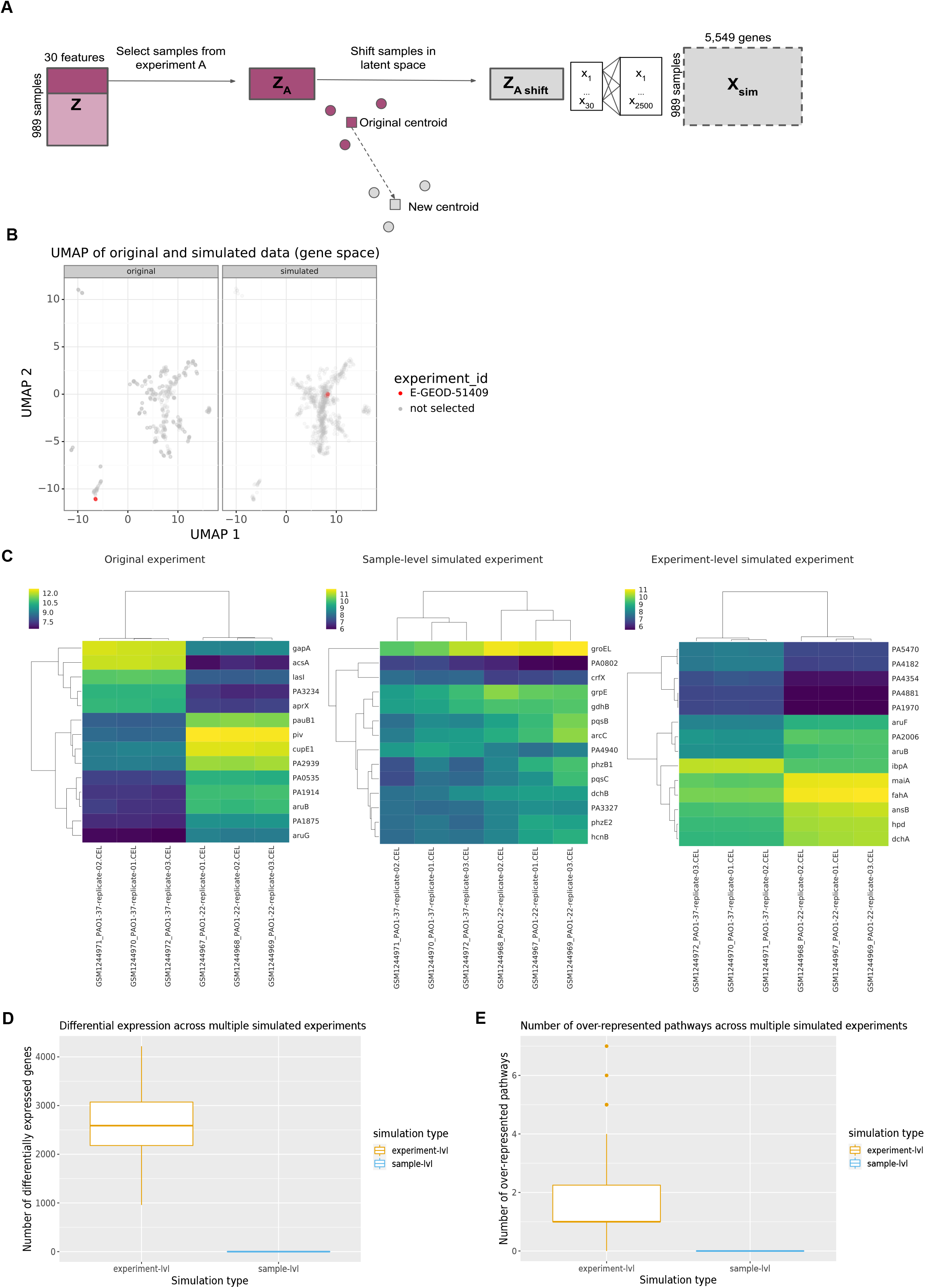
Simulating gene expression compendia by experiment. A) Workflow to simulate gene expression per experiment. B) UMAP projection of P. aeruginosa gene expression data highlighting a single experiment, E-GEOD-51409, (red) in the original dataset (left) and the simulated dataset (right), which was subsampled to 1000 samples. C) Differential expression analysis of experiment E-GEOD-51409 (left), random simulated samples (middle), simulated samples using the same experiment as a template (right). D) Number of differentially expressed genes identified across 100 simulated experiments generated using experiment-level simulation and sample-level simulation. E) Number of enriched pathways identified across 100 simulated experiments generated using experiment-level simulation and sample-level simulation.

Given the fact that we preserved the association between samples and experiments in this new simulation, we would expect that new experiments would preserve the correlation in expression of genes that are in the same pathway. In our previous example experiment, E-GEOD-51409, the simulated experiment generated using the original E-GEOD-51409 as a template (i.e. experiment-level) identified 14 differentially expressed genes (Figure 3C). In contrast, the simulated experiment generated by randomly sampling (i.e. sample-level) did not identify any differentially expressed genes; the median log2 fold change was 0.08. Furthermore, when simulating 100 new experiments using E-GEOD-51409 as a template, the experiments generated using the workflow in Figure 3A identified a median of 2,588 differentially expressed genes compared to those new experiments generated by randomly sampling from the compendium resulting from the workflow in Figure 1C (Figure 3D) which identified a median of 0 differentially expressed genes. Additionally, the median number of enriched KEGG pathways is 1 using the workflow in Figure 3A compared to 0 using the random sampling approach using the previous simulation strategy (Figure 3E). Overall, it appears that this new simulation approach generated a compendium of experiments with some real underlying biology and therefore this new simulation represents a more realistic simulation compared to the previous one. Examples of the significantly enriched pathways can be seen in Table 2. The top over-represented pathway is the ribosome pathway, which is likely a commonly altered pathway found in many experiments regardless of experiment type.^26^ The remaining pathways found in the original experiment were generally metabolism related, which is consistent with the finding from the original publication.^25^ The simulated experiment was particularly enriched in sulfur metabolism and ABC transporters, which is consistent with a different previous experiment that found upregulation of transport systems in response to sulfate limitations.^27^ Overall, in accordance with real gene expression experiments, the new simulated experiments contain related groups of enriched pathways which reflect the specific hypothesis being tested.

**Table 2:** Enriched pathways found in the original E-GEOD-51409 experiment and the pseudo-experiment generated using the experiment-level simulation.

| Original | Adjusted p-value | Experiment level simulation | Adjusted p-value |
| --- | --- | --- | --- |
| Pae03010: Ribosome | 2.966E-11 | Pae03010: Ribosome | 7.96E-07 |
| Pae00500: Starch and sucrose metabolism | 0.001512 | Pae02010: ABC transporters | 0.004009 |
| Pae01200: Carbon metabolism | 0.004466 | Pae00920: Sulfur metabolism | 0.01576 |
| Pae00640: Propanoate metabolism | 0.001954 |  |  |

### Simulating gene expression compendia with synthetic experiments

We used our method to simulate new experiments that follow existing patterns to examine the patterns that we observed for generic partitions (Figure 4A). We simulated 600 experiments using the *P. aeruginosa* compendium. We divided these experiments into partitions. These partitions represent groupings of experiments with shared noise, such as experiments from the same lab or experiments with the same experimental design. Each partition contains technical sources of variance within and between experiments. Results with simulated experiments were similar to those from arbitrarily partitioned samples. We observed a monotonic loss of similarity after the correction step as the number of partitions increased (Figure 4B). Visualizing the top principal components revealed that statistical correction initially better recapitulated the overall structure of the data but that with many partitions similarity decreased (Figure 4C, dark blue). Without correction there was a larger initial drop in similarity but a later recovery (Figure 4B) and visualizing the top principal components recapitulated this finding (Figure 4C, light blue). We performed analogous experiments using the recount2 VAE and 100 simulated experiments. We observed consistent results with this dataset using both SVCCA similarity (Figure 4D) and visual inspection of the top principal components (Figure 4E). In summary, as the number of partitions increase the experiment-specific technical sources contribute less overall to the signal and the underlying patterns dominate the overall signal. When many partitions are present, even ideal statistical approaches to correct for noise over-corrects and removes the underlying signal.

**Figure 4.**
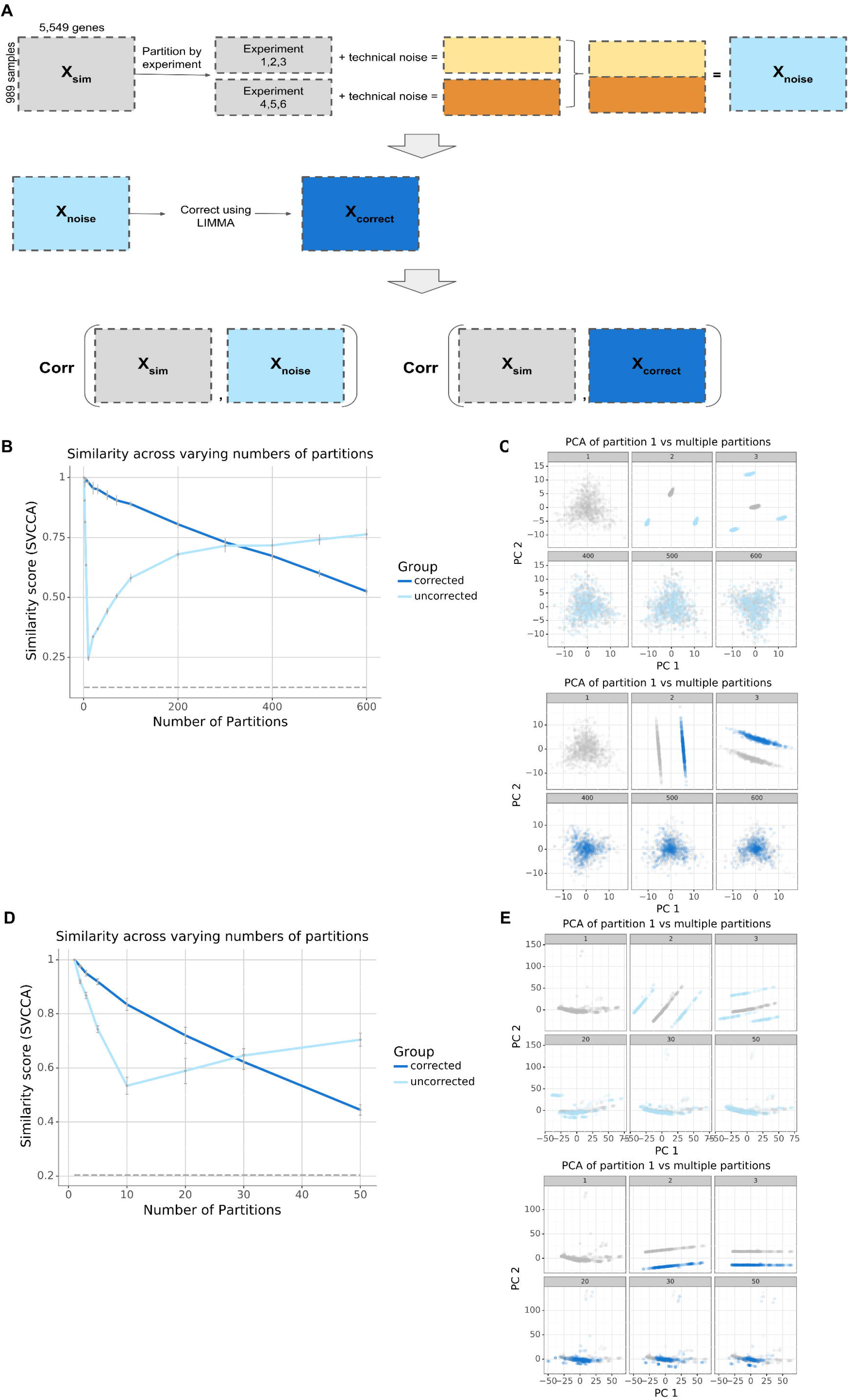
Results of simulating compendia comprised of gene expression experiments. A) workflow describing how experiment-specific noise was added to the simulated compendia and how the simulated compendia were evaluated for similarity compared to the original input compendia. B,D) SVCCA curve measuring the similarity between a compendia without noise versus a compendium with noise (light blue), compendium with noise corrected for (dark blue). As a negative control, we used the similarity between the gene expression pattern of the simulated data with a single partition compared with the simulated data that has been permuted to destroy any meaningful structure in the data. C,E) Subsampled gene expression data (500 samples per compendia) projected onto the first two principal components showing the overlap in structure between the compendia without noise (gray) versus the compendia with noise (light blue), compendia with noise corrected for (dark blue).

## Discussion

Our findings reveal that compendia-wide analyses do not always require correction for experiment-specific technical variance and that correcting for such variance may remove signal. This simulation study provides an explanation for the observation that past studies^2–6^ have successfully extracted biological signatures from gene expression compendia despite the presence of uncorrected experiment-specific sources of technical variability. In general, there exists compendia that contain some small number of experiment-specific sources where traditional correction methods can be effective at recovering the biological structure of interest; however, there also exist large-scale gene expression compendia where these methods may be harmful instead of helpful. The number of experiment-specific sources that determine whether to apply correction will vary depending on the dataset – the size of the compendia, the magnitude and structure of the signals. Using the associated repository (https://github.com/greenelab/simulate-expression-compendia) users can customize the scripts to run the simulation experiments on their own expression data in order to examine the effect of a linear noise model on their dataset. Though our analysis uses simplifying assumptions that preclude us from defining a specific threshold for noise correction, these simulations define a set of general properties that will guide compendia analyses moving forward.

We introduce a new method to simulate genome-wide gene expression experiments, using existing gene expression data as starting material, which goes beyond simulating individual samples. This allowed us to examine the extent to which our findings hold with realistic experimental designs. The ability to simulate gene expression experiments with a realistic structure may have many potential legitimate uses: e.g., pre-training for machine learning models, providing synthetic test data for software, and other such applications. Additionally, this simulation technique can be used to explore hypothetical experiments that have not been previously performed and generate hypotheses. However, such approaches could also be used by nefarious actors to generate synthetic data for publications. Forensic tools that detect synthetic genome-wide data may be needed to combat potential fraudulent uses.

Our study has several limitations. We assume a certain noise model that differs between experiments. However, the sources of noise are multifaceted and any such assumption will necessarily be an oversimplification, though such assumptions are not uncommon.^10,12,28^ By selecting a specific noise model and using an ideal noise-removal step, we provide a best case scenario for artifact removal. While any simulation study will necessarily make simplifying assumptions, this work is the first to use deep generative models as part of a simulation study to probe the long-standing assumption that correcting for technical variability is necessary for analyses that span multiple experiments. Our findings reveal that in settings with hundreds or thousands of experiments, correcting for experiment-specific effects can harm performance and that it can be best to do nothing.

Our study also has broader implications for efforts to standardize scientific processes. Centralization of large-scale data generation has the potential to reduce experiment-specific technical noise, though it comes at a cost of flexibility. Our results suggest that a highly distributed process where experiments are carried out in many different locations with their own specific sources of technical noise can also lead to valuable data collections.

## Methods

### Pseudomonas aeruginosa gene expression compendium

We downloaded a compendium of *P. aeruginosa* data that was previously used for compendium-wide analyses.^2^ Previous studies identified biologically-relevant processes such as oxygen deprivation^2^ and phosphate starvation^3^ by applying denoising autoencoders. We obtained the processed and normalized gene expression matrices from the associated GitHub repository. The *P. aeruginosa* dataset was previously processed by Tan et. al.^2^ During processing, raw microarray data were downloaded as .cel files, *rma* was used to convert probe intensity values from the *.cel* files to log2 base gene expression measurements, and these gene expression values were then normalized to 0-1 range per genes.

This compendium includes measurements from 107 experiments that contain 989 samples for 5,549 genes.^2^ It contains experiments that accrued between the release of the GeneChip *P. aeruginosa* genome array and at the time of data freeze in 2014. Approximately 70% of the samples were from cultures of strain PAO1 and derivatives, 13% were in strain PA14 background, 0.6% were from PAK strains and the remaining were largely clinical isolates. Of the strains, 73% were wild-type (WT) genotypes and the rest were mutants that had undergone genetic modification. Approximately 60% of the samples were grown in LB medium while the rest were grown in Pseudomonas Isolation Agar (PIA), glucose, pyruvate or amino acid-based media.^3^ Roughly 80% were grown planktonically, 15% were grown in biofilms and the remaining samples were in vivo or not annotated. Overall, this *P. aeruginosa* compendium covered a wide range of gene expression patterns including characterization of clinical isolates from Cystic Fibrosis infections, response of mutant versus WT, response of antibiotic treatment, microbial interactions, adaptation from water to GI tract infection. Despite having 989 samples, this compendium represents the heterogeneity of *P. aeruginosa*.

### recount2 gene expression compendium

We downloaded human RNA-seq data from recount2.^29^ The dataset includes over 70,000 samples collected from Sequencing Read Archive (SRA). It is comprised of more than 50,000 samples from different types of experiments, roughly 10,000 samples from Genotype-Tissue Expression project (GTEx v6) covering 44 types of normal tissue, and more than 10,000 samples from The Cancer Genome Atlas (TCGA) measuring 33 cancer types.^19,30,31^ The recount2 authors uniformly processed and quantified these data. We downloaded data using the recount library in Bioconductor (version 1.14.0).^29^ The entire recount2 dataset is 8TB. Based on the *P. aeruginosa* compendium we expected that a subset of the compendium would be sufficient for this simulation, so we selected a random subset of 50 NCBI studies, which resulted in 896 samples with 58,037 genes for our simulation. Each project (imported from NCBI BioProject) is akin to an experiment in the *P. aeruginosa* compendium, and we used the term *experiment* instead of projects in order to maintain consistency in this paper. The downloaded recount2 dataset was in the form of raw read counts, which was normalized to produce RPKMs used in our analysis. The normalized gene expression data was then scaled to a 0-1 range per gene.

### Constructing a generative model of gene expression compendia

We designed an approach to simulate gene expression compendia with a multi-layer variational autoencoder (VAE). We built this model in Keras (version 2.1.6) with a TensorFlow backend (version 1.10.0) based on the Tybalt software for gene expression VAEs.^32–34^ Our architecture used each input gene as a feature. We compressed these genes to 2,500 intermediate features using a rectified linear unit (ReLU) activation function to combine weighted nodes from the previous layer. We encoded these features into 30 latent space features, which we optimized to follow a standard normal distribution via the addition of a Kullbach-Leibler (KL) divergence term into the loss function. We then reconstructed these features back to the input space using decoding layers that mirror the encoder network. We trained the VAE using 90% of the input dataset, leaving 10% as a validation set. We determined training hyperparameters by manually adjusting parameters and selecting parameters that optimized the validation loss. These were a learning rate of 0.001, a batch size of 100, warmups set to 0.01, 100 epochs for the *P. aeruginosa* compendium and 20 epochs for the recount2 compendium.

We used the VAE trained from each compendium to generate new compendia by randomly sampling from the latent space. We generated a simulated compendium containing 6,000 *P. aeruginosa* samples or 500 recount2 samples. For our first simulation, we sampled randomly - ignoring the relationship between samples with a specific experiment. We simulated experiment-specific sources of undesired variability within compendia by dividing the data into partitions and adding noise to each partition.

We divided the *P. aeruginosa* simulated compendium into [1, 2, 5, 10, 20, 50, 100, 500, 1000, 2000, 3000, 6000] partitions and divided the recount2 simulated compendium into [1, 2, 5, 10, 20, 50, 100, 250, 500] partitions. We randomly added linear noise to each partition by generating a vector of length equal to the number of genes (5,549 *P. aeruginosa* genes and 58,037 Human genes) where each value in the vector was drawn from a normal distribution with a mean of 0 and a variance of 0.2. With the 0-1 scaling, a value of 0.2 produced a relatively large difference in gene expression space (Figure S1), which allowed us to evaluate the impact of a large amount of technical noise.

Though linear noise is an over-simplification of the types of noise that affect gene expression data, it allowed us to design an approach to optimally remove noise. Adjusting the choices of normalization, noise magnitude, and noise patterns will result in different selections of the precise cross-over point where it becomes beneficial to perform correction. With this design, we do not expect that it is possible to estimate exactly where this precise cross-over point is. That would require a compendium where investigators systematically performed the same combination of different experiments in multiple labs at different times. We were unable to identify such a compendium on the scale of thousands of samples from tens to hundreds of labs. Thus, though our analysis necessarily includes simplifying assumptions that limit our ability to precisely define the thresholds for correction for arbitrary datasets and noise sources, it remains suitable for examining the overriding principles that govern compendium-wide analyses.

We also designed an approach to generate gene expression compendia with realistic experimental designs. There was no consistent set of annotated experimental designs, so we developed a simulation method that did not depend on *a priori* knowledge of experimental design. For each synthetic experiment, we randomly sampled a “template experiment” from the set of *P. aeruginosa* or recount2 experiments. We then simulated new data that matched the template experiment by selecting a random sample from the low dimensional representation of the simulated compendia and calculating the vector (Z_shift_) that connected this random sample (Z_random_) and the encoded template experiment (X_template_experiment_) using equation 1:

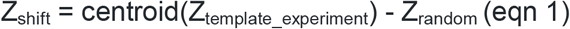

We then linearly shifted the template experiment in the low-dimensional latent space by adding this vector to each sample in the experiment as seen in equation 2.

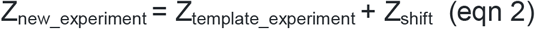

This process preserves the relationship between samples within the experiment but shifts the samples to a new location in the latent space. Intuitively this simulation maintains the same experimental design but is akin to studying a distinct biological process. Repeating this process for each experiment allowed us to generate new simulated compendia comprised of realistic experimental designs.

We divided the *P. aeruginosa* simulated compendium into [1, 2, 3, 5, 10, 20, 30, 50, 70, 100, 200, 300, 400, 500, 600] partitions and divided the recount2 simulated compendium into [1, 2, 5, 10, 20, 30, 50] partitions, where experiments are divided equally amongst the partitions. For each partition we added simulated noise as described in the previous section. Experiments within the same partition have the same noise added. Each partition represents a group of experiments generated from the same lab or with the same experimental design.

### Removing technical variability from noisy compendia

Our model of undesired variability is a linear signature applied separately to each partition of the data, which we consider akin to experiments or groups of experiments in a compendium of gene expression data. We used the removeBatchEffect function in the R library, limma (version 3.44.0), to correct for the technical variation that was artificially added to the simulated compendia.^24^ Limma removes the technical noise by first fitting a linear model to describe the relationship between the input gene expression data and the experiment labels. The input expression data contains both a biological signal and technical noise component. By fitting a linear model, limma will extract the noise contribution and then subtract this from the total input expression data. This method presents a best-case scenario for removing the undesired variability in the simulated compendia because the model matches the noise pattern we’ve used in the simulation.

### Measuring the similarity of matched compendia

We used Singular Vector Canonical Correlation Analysis (SVCCA)^35^ to estimate similarities between different compendia. SVCCA is a method designed to compare two data representations^35^. Given two multivariate datasets, X_1_ and X_2_, the goal of SVCCA is to find the basis vectors, *w* and *s*, to maximize the correlation between w^T^X^1^ and s^T^X_2_. In other words, SVCCA attempts to find the space, defined by a set of basis vectors, such that the projection of the data onto that space is most correlated. Two datasets are considered similar if their linearly invariant correlation is high (i.e., if X_1_ is a shift or rotation of X_2_ then X_1_ and X_2_ are considered similar).

We compared the statistical structure of the gene expression, projected onto the first 10 principle components, in the baseline simulated compendia (those with only one experiment or partition, X_1_) versus those with multiple experiments or partitions (X_2_). Our SVCCA analysis was designed to measure the extent to which the gene expression structure of the compendia without noise is similar to the gene expression structure of the compendia with multiple sources of technical variance has been added as well as those where correction has been applied. Here we use 10 principle components for computational simplicity. Selecting a different value would affect the crossover point but not the general trends that we describe.

### A case study of differential expression in a template experiment

We compared the E-GEOD-51409 experiment^36^ with two different simulated representations to provide a case study for experiment-based simulation. E-GEOD-51409 included *P. aeruginosa* in two different growth conditions. For one simulation, we generated random samples and randomly assigned them to conditions, which we termed the sample-simulated experiment. For the second we used the latent space transformation process described above, which we termed the experiment-simulated experiment. We used the eBayes module in the limma library to calculate differential gene expression values for each gene between the two different growth conditions in the real and simulated data. We built heatmaps for the 14 most differentially expressed genes, where differentially expressed genes where those with FDR adjusted cutoff < 0.05 and log2 fold change >1, which are thresholds frequently used in practice. We selected 14 genes because there were 505, 14, 0 differentially expressed genes found in the original experiment, experiment-simulated experiment, sample-simulated experiment respectively. Since there were 0 differentially expressed genes found in the sample-simulated experiment, we displayed the top 14 genes sorted by adjusted p-value to provide a visual summary of the simulation process.

### Comparing sample-level and experiment-level simulated datasets

We simulated 100 experiments using the template E-GEOD-51409 experiment^36^. We sought to compare the sample-level and experiment-level simulation processes. We set a threshold for differentially expressed genes at a Bonferroni-corrected p-value cutoff of 0.05. We used the enrichKEGG module in the clusterProfiler library to conduct an over-representation analysis^37^. We used the Fisher’s exact test to calculate a p-value for over-representation of pathways in the set of differentially expressed genes. We considered pathways to be over-represented if the Bonferroni corrected p-value was less than 0.05.

### Implementation and Software Availability

All scripts to reproduce this analysis are available the GitHub repository (https://github.com/greenelab/simulate-expression-compendia) under an open source license. The repository contains 98% python jupyter notebooks, 2% python and 0.1% R scripts. The repository’s structure is separated by input dataset. Pseudomonas/ and Human/ directories each contain the input data in the data/input/ directory. Scripts for the sample level simulation can be found in Pseudomonas /Pseudomonas_sample_lvl_sim.ipynb for the *P. aeruginosa* compendium and Human/Human_sample_lvl_sim.ipynb for the recount2 compendium. Scripts for the experiment level simulation can be found in Pseudomonas/Pseudomonas_experiment_lvl_sim.ipynb and Human/Human_experiment_lvl_sim.ipynb respectively. The virtual environment was managed using conda (version 4.6.12), and the required libraries and packages are defined in the environment.yml file. We describe in the Readme file how users can analyze different compendia or use different noise patterns.

---

**Figure S1**. Results of varying the magnitude of the experiment-specific noise added to each partition. SVCCA curve measuring the similarity between a compendium without noise versus a compendium with noise (light blue), compendium with noise corrected for (dark blue). As a negative control, we used the similarity between the gene expression pattern of the simulated data with a single partition compared with the simulated data that has been permuted to destroy any meaningful structure in the data. Using noise model with A) 0.2 variance, B) 0.05 variance with a zoomed in view on the left, C) 0.025 variance with a zoomed in view on the left.

**Figure S2**. Simulating recount2 gene expression data using VAE. A) Validation loss plotted per epoch during training. B) UMAP projection of gene expression data from the real dataset (pink) and the simulated compendium using the workflow in Figure 1C (grey).

**Figure S3**. Results of simulating *P. aeruginosa* compendia using A) sample-level simulation or B) experiment-level simulation with COMBAT noise correction.

**Figure S4**. Clustering of 100 random gene expression profiles in original versus simulated experiments using A) E-GEOD-21704 and B) E-GEOD-10030 as templated.

## Notes

https://github.com/greenelab/simulate-expression-compendia

### Competing Interest Statement

The authors have declared no competing interest.

