## Supplementary figures and images for "Correcting for experiment-specific variability in expression compendia can remove underlying signals"

### Figure S1

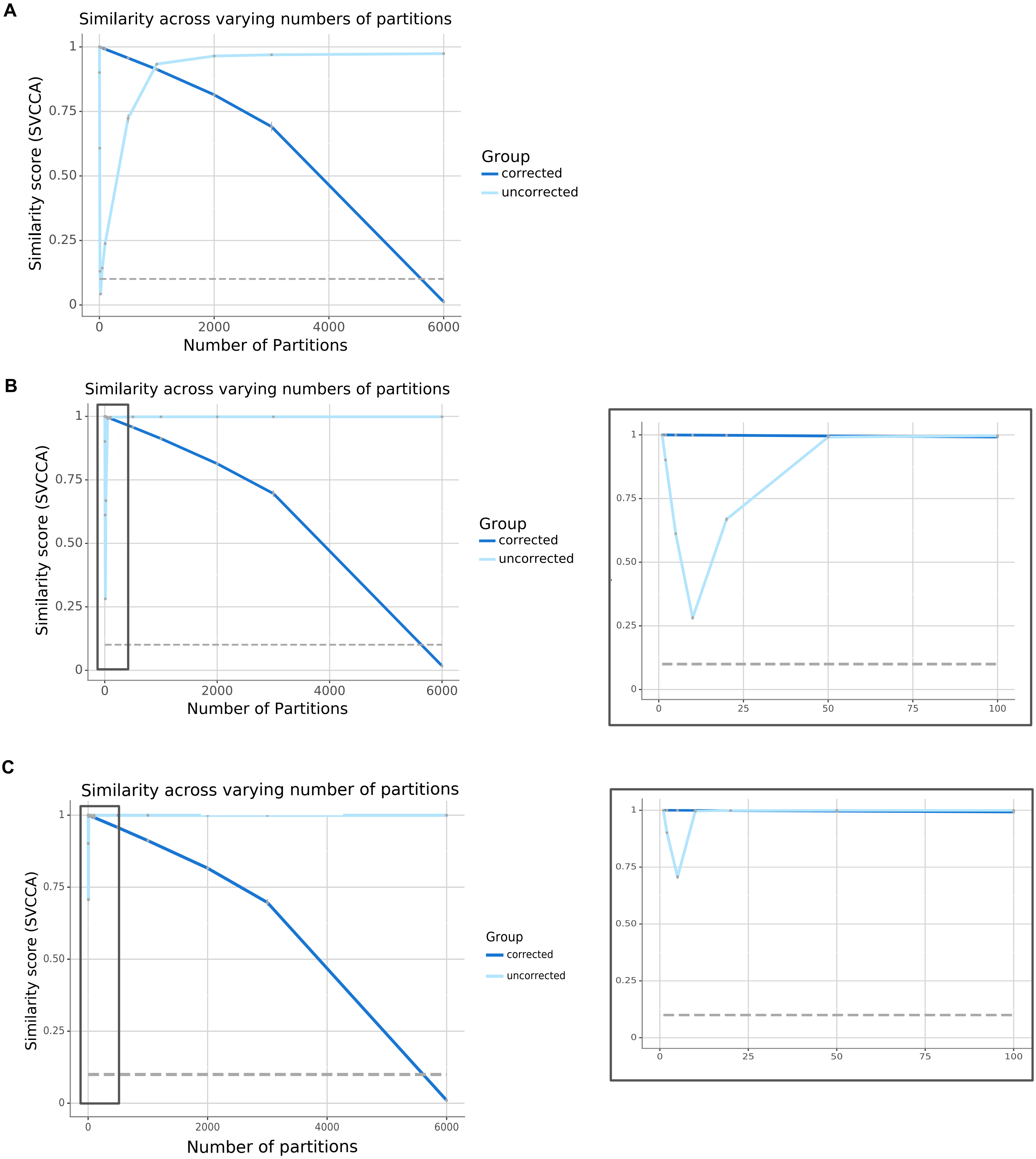

### Figure S2

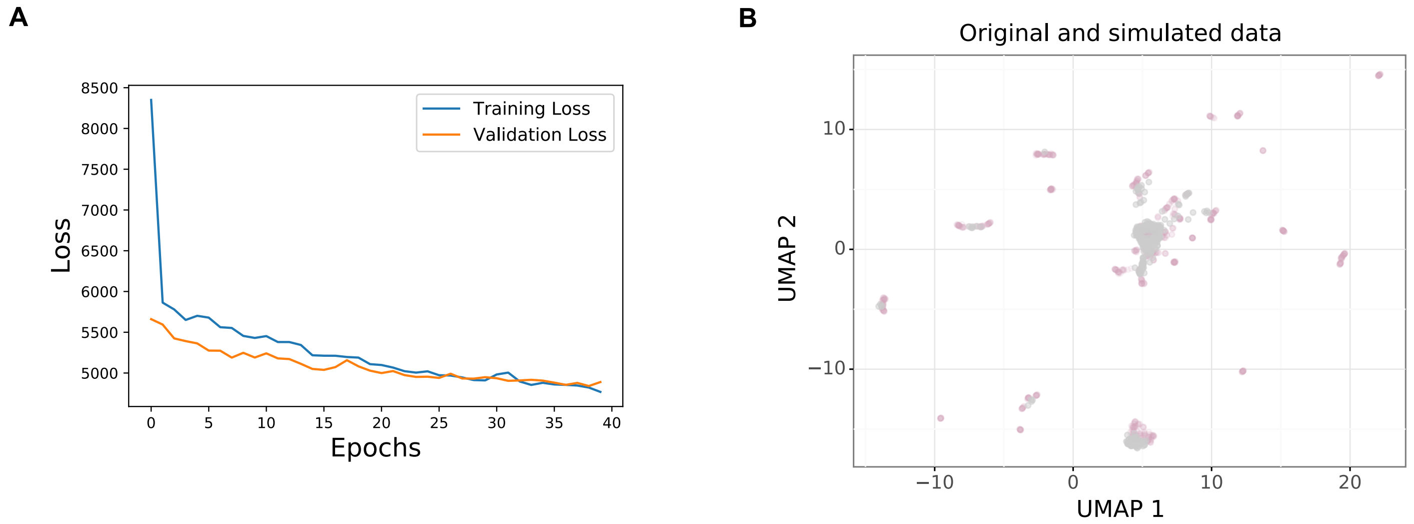

### Figure S3

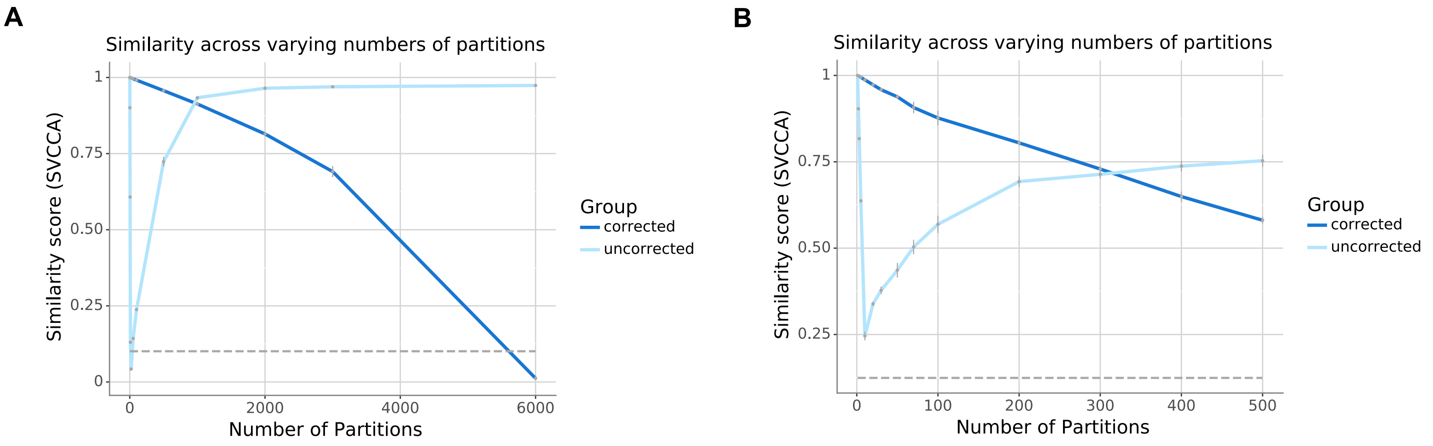

### Figure S4

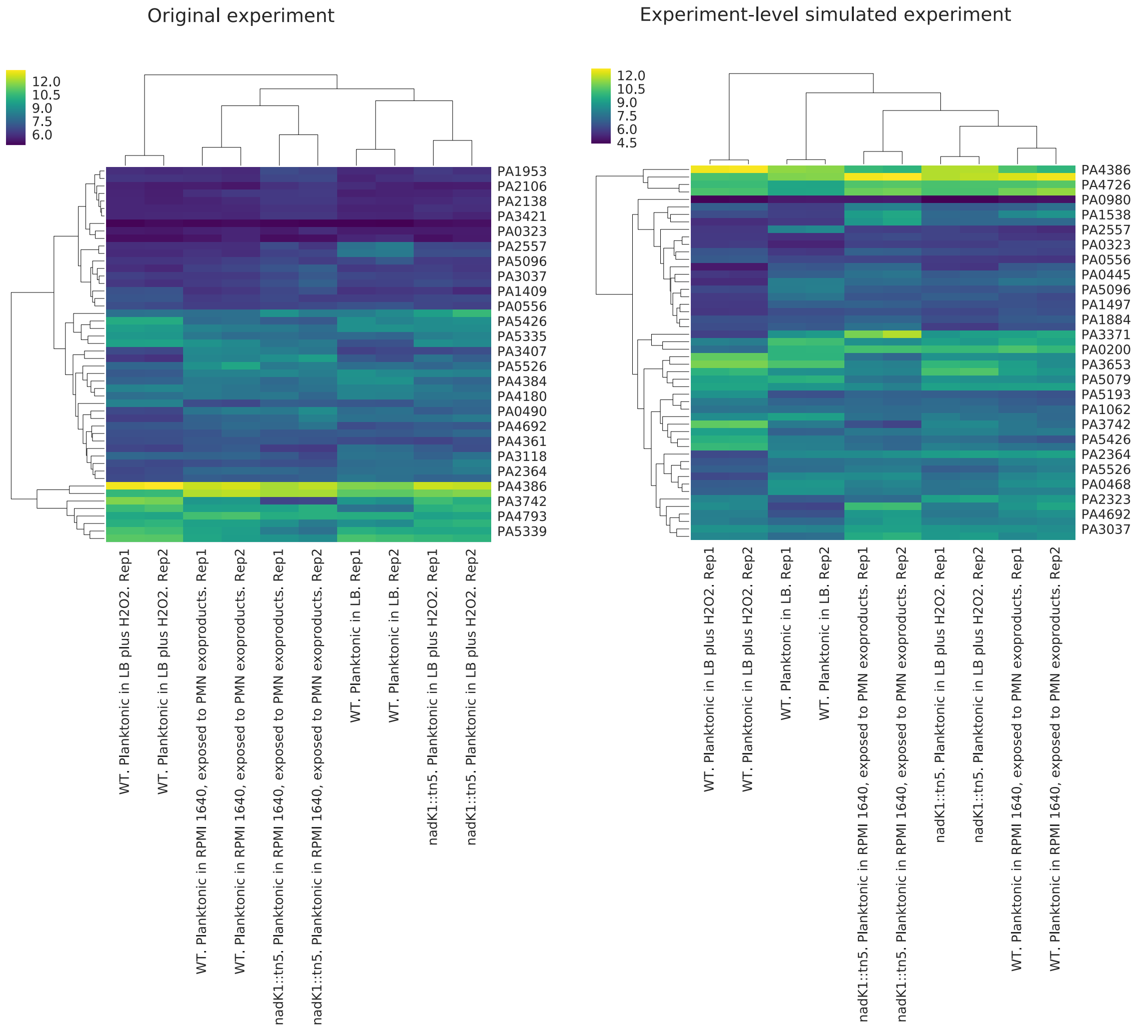

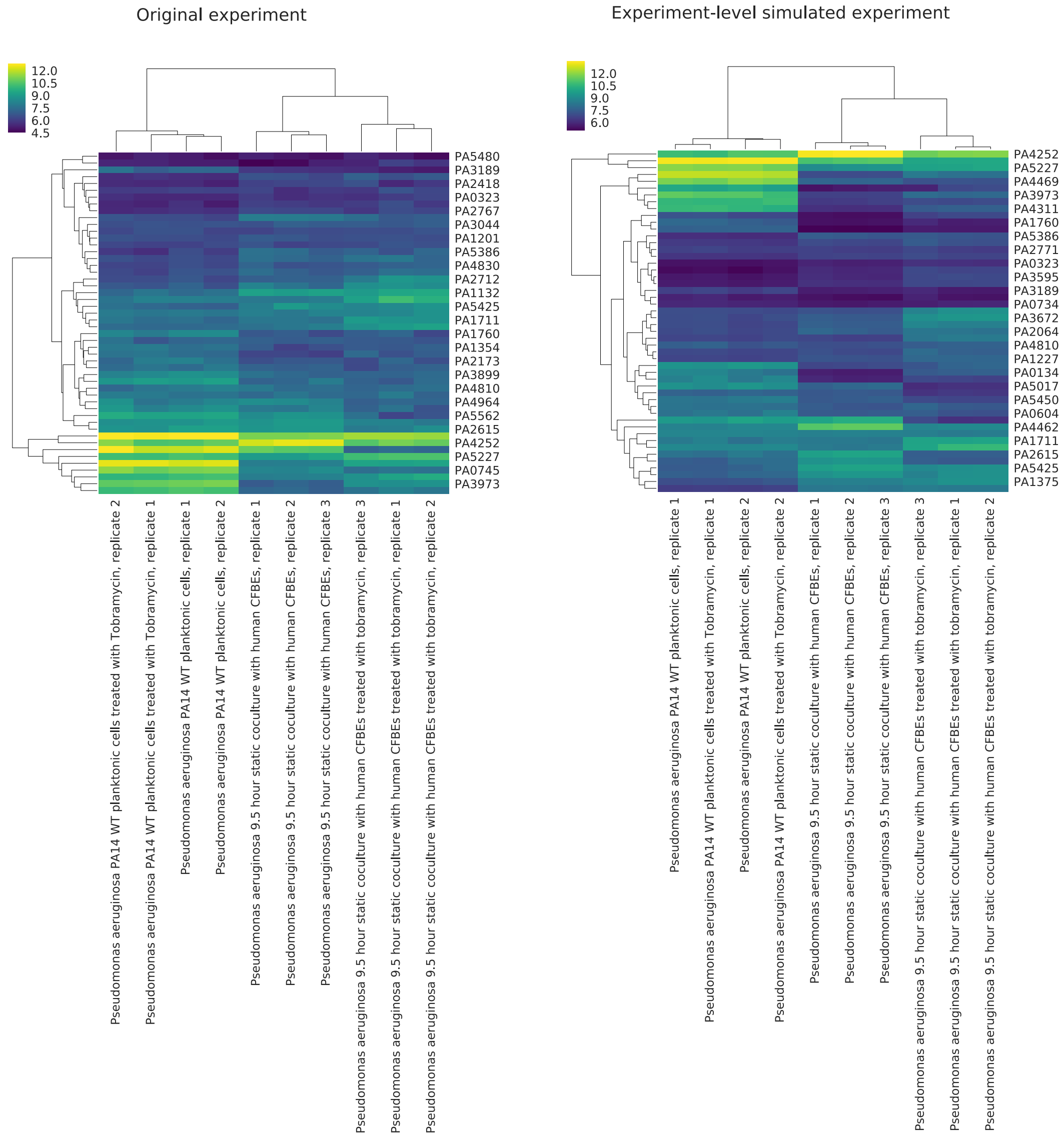
